# Detritus identification in FlowCAM using a simple binary classifier

**DOI:** 10.1101/2024.11.18.624123

**Authors:** O. García-Oliva

## Abstract

Phytoplankton and detritus particles may be co-captured in Flow-CAM systems, leading to misrepresentation of phytoplankton abundance. In this study, I use logistic regression as a binary classifier to identify detritus particles in FlowCAM data from the Southern North Sea. Standard particle features from the manufacturer’s software were used as inputs, with surface texture (intensity variance) and compactness (derived from particle perimeter and area) proving most relevant to detritus classification. This classifier achieved 81% accuracy using a training dataset of approximately 7300 observations, reducing the workload compared to other classification methods. The reconstructed particle size spectra closely matched the observed spectra for detritus and phytoplankton. Binary classifiers like this offer a fast, effective alternative for detritus screening, aiding the pre-processing and re-analysis of FlowCAM datasets.

## 1 Introduction

Analysing phytoplankton size-spectra can reveal key ecosystem processes, making accurate size-spectrum description essential [20, 6]. Under suitable conditions, FlowCAM analyses can reconstruct phytoplankton size spectra, often with assistance from machine learning algorithms that classify particles into phytoplankton classes [2]. However, high concentrations of detritus particles can lead to overestimating phytoplankton, impeding the accurate acquisition of size-spectra [7, 9, 2]. Classifying detritus particles within FlowCAM samples usually involves image analysis and classification algorithms [2]. The complexity of these algorithms varies, with simpler classifiers used in controlled settings and more complex schemes for natural samples and higher taxonomic classifications [1, 2, 4, 12, 16, 18, 17]. Here I apply a simple logistic regression classifier to identify detritus particles in Southern North Sea samples, evaluating its effectiveness in reconstructing detritus and phytoplankton size spectra of natural samples.

## 2 Material and methods

### 2.1 Data management

The dataset used for this study consists of particle samples collected in the Southern North Sea from November and December 2007 [9, 6]. Observations were captured using a portable black-and-white FlowCAM system in fluorescence-triggered image mode with settings recommended by the manufacturer [5, 9]. The dataset includes particle classifications as either detritus or phytoplankton taxa [9, 6], along with standard particle variables provided by the software Visual SpreadSheet version 1.5.16 [10, 5]. I divided these variables into four groups: (1) dimensional –sum intensity, volume, area, diameter, convex perimeter, and perimeter–, (2) shape –elongation, compactness, and aspect ratio–, (3) texture –intensity, sigma intensity, and roughness–, and (4) optical –transparency, fluorescence peak Ch. 1, and fluorescence peak Ch. 2. Table 1 contains descriptions of these variables. Incomplete records (i.e., those with missing or invalid values) were removed from the dataset.

**Table 1:** List of particle features provided by Visual SpreadSheet software and used herein (symbol in parenthesis) [5, 17].

| Feature | Description |
| --- | --- |
| **Dimensional** |  |
| Volume (VOL) | Sphere volume calculated from the diameter (ESD). |
| Area (AREA) | Number of pixels in the thresholded (binary) greyscale image converted to a measure of the area by use of the calibration factor. |
| Diameter (ESD) | The mean value of 36 ferret measurement, from these the larger value is the **Length** and the smaller as the **Width** of the particle. |
| Convex perimeter (CONP) | An approximation of the perimeter of the convex hull of a particle. |
| Perimeter (PER) | Total length of the edges making up a particle including the edges of any holes. Perimeter is calculated from the **Length** and **Breadth** of the particle by $PER = 2 \times (Length + Breadth)$. |
| Sum intensity (SUMIN) | Sum of greyscale pixel values. |
| **Shape** |  |
| Elongation (ELON) | A **Length** over **Breadth** ratio based on PER and AREA with the assumption that $AREA = Length \times Breadth$. 1 is the value for a filled circle or square; larger values are for elongated particles. |
| Compactness (COMP) | A shape parameter derived from the PER and AREA. |
| Aspect ratio (AR) | **Width** over **Length**. 1 is the value for a perfect circle; values near zero are for particles that are long and thin. |
| **Texture** |  |
| Intensity (INT) | The average greyscale value of the pixels making up a particle. |
| Sigma intensity (SIGMAI) | Standard deviation of greyscale values. |
| Roughness (ROUGHN) | A measure of the unevenness or irregularity of a particle's surface—the ratio of perimeter to convex perimeter. 1 is the value for a filled shape with a convex perimeter; larger values are for particles with interior holes or a non-convex perimeter. |
| **Optical** |  |
| Transparency (TRANS) | Calculated from the ratio of the diameter derived from AREA—the Area Based Diameter, ABD— and ESD by $TRANS = 1 - ABD/ESD$. 0 is the value for a filled circle, values near 1 are for an elongated or irregular shape or a shape that has many interior holes. |
| Ch.1 peak (CH1) | Peak fluorescence value read from PMT #1 (>650 nm filter). Red fluorescence is associated with chlorophyll content. |
| Ch.2 peak (CH2) | Peak fluorescence value read from PMT #2 (575 nm filter). |

### 2.2 Logistic models as a simple binary classifier

Logistic regression can serve as a binary classifier for data with a binary outcome variable, where the target follows a Bernoulli distribution, i.e., a variable taking the value 1 with probability *p* and 0 with probability 1 − *p* [11]. Here, *p* represents the probability of a particle being detritus. Logistic models assume that the log odds of *p* are a linear function of the particle features:

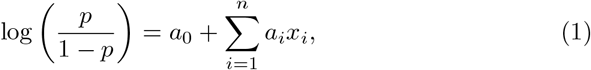

where *x*_*i*_ represents normalized particle features, and *a*_*i*_ are regression coefficients indicating the relative importance of each variable for detritus classification. Only non-dimensional variables were used to fit the logistic model, ignoring size-related biases in particle composition.

The model was fitted 1000 times using randomly selected training datasets (30% of the valid observations). Model performance was evaluated on the remaining test dataset by comparing predicted and observed classifications. Evaluation metrics included accuracy, precision, recall, and F1-score, following other classification schemes [e.g., 2, 14, 17]. Score metrics and regression coefficients were also calculated using a random selection ranging from 1–30% of the observations from the dataset, iterated 1000-fold to calculate uncertainties as standard deviation. These metrics were based on the confusion matrix (see SM A for definitions). The percentage of misidentified particles per date and size-class (<2, 2-5, 5-10, 10-20, 20-50, 50-100, and 100-200 μm intervals) were also reported.

All analyses were performed using Python’s scikit-learn package https://scikit-learn.org/1.5/index.html, with code available at https://github.com/ovgarol/detritus-logistic-classification.

## 3 Results

The dataset included a total of 86,930 observations. After removing incomplete records, the dataset was reduced to 24,598 valid observations. From these, 13,145 (53.4%) records were detritus particles and 11,456 (46.6%) were phytoplankton. Using a training dataset of 30% of the valid observations, the logistic classifier achieves: Accuracy = 0.8115 ±0.0017, Precision = 0.8249 ±0.0042, Recall = 0.8219 ±0.0053, and F1-score = 0.8234 ±0.0017. These performance metrics remained nearly constant when the size of the training set was comprised of more than 10% of the total observations, with marginal improvements between in the range 1-10% (Fig. 1).

**Figure 1.**
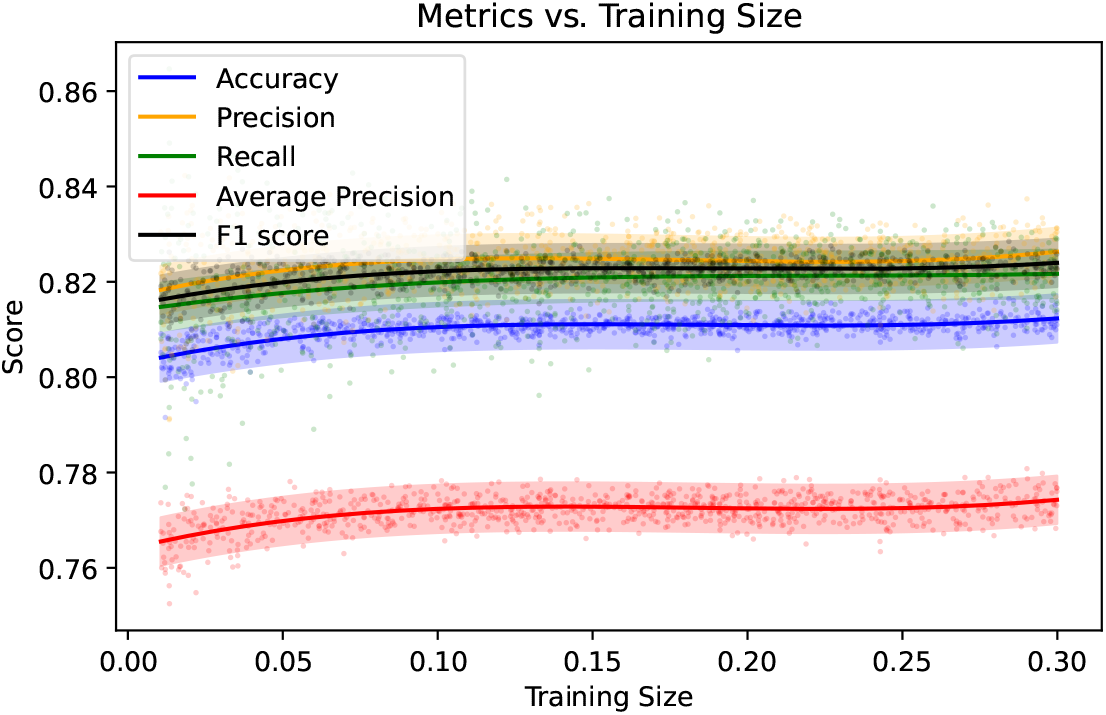
Evaluation metrics as a function of the size of the training dataset.

Features related to surface texture –sigma intensity– and shape –compactness– appear as determinant variables for detritus identification during model fitting (Table 2). Sigma intensity and compactness importances are similar and followed by other shape and texture features, such as elongation, intensity and roughness. Optical properties are the least important in the identification of detritus particles.

**Table 2:** Particle features sorted by absolute importance, mean and standard deviation (SD).

| Feature | Mean $a_i$ | SD |
| --- | --- | --- |
| SIGMAI | 2.264 | 0.119 |
| COMPACT | -2.069 | 0.107 |
| ELON | 1.345 | 0.145 |
| INT | -1.261 | 0.108 |
| ROUGHN | 1.254 | 0.082 |
| CH1 | -0.799 | 0.032 |
| CH2 | -0.358 | 0.030 |
| AR | -0.277 | 0.054 |
| TRANS | 0.236 | 0.059 |

The predicted and observed detritus and phytoplankton particle abundance size-spectra match along the equivalent spherical diameter range from 1 to 100 μm (Fig. 2). In general, our classifier slightly underestimated detritus particle abundances in the 2-5 and 5-10 μm size classes. Overestimation occurs more frequently and with higher percentages in the 10-20 μm size class. Larger particles appear more difficult to classify (Table 3).

**Table 3:** Particle misidentification per size-class and date.

| size-class ($\mu\text{m}$) | <2 | 2-5 | 5-10 | 10-20 | 20-50 | 50-100 | 100-200 |
| --- | --- | --- | --- | --- | --- | --- | --- |
| 20071115 | – | – | – | 1.4 | -5.6 | – | – |
| 20071122 | 0.0 | 7.6 | -0.7 | -8.6 | -22.6 | -35.0 | 50.0 |
| 20071129 | 0.0 | -1.6 | 6.6 | 1.3 | 1.0 | 6.6 | 100.0 |
| 20071206 | 0.0 | 3.6 | 7.4 | 11.2 | 6.8 | 9.0 | – |
| 20071213 | -2.7 | -3.7 | -3.5 | -0.7 | -1.1 | -28.5 | -100.0 |
| 20071220 | 2.0 | 4.9 | -2.2 | -4.1 | -7.6 | 5.2 | – |

**Figure 2.**
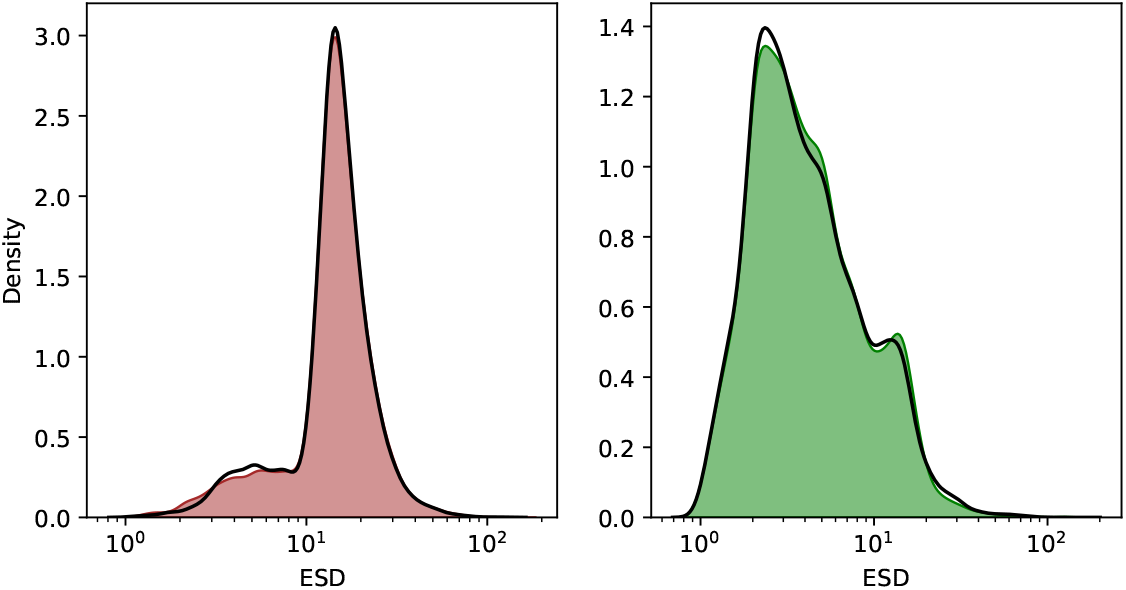
Observed (black line) and predicted abundance size spectra (ESD, Equivalent Spherical diameter in μm) for detritus (brown area) and phytoplankton (green area).

## 4 Discussion

The abundant data captured by FlowCAM systems motivates the use of automated and semi-automated classification techniques to replace the otherwise laborious manual work [2]. Machine learning is used for micro-algae classification using image processing in taxa identification [18, 17]. Sophisticated deep learning techniques provide accuracies greater than 95% [14] in particle identification. In comparison, simpler methods report accuracies of 80–90% [4, 2], as calculated herein (Fig. 1).

FlowCAM-based monitoring programs can achieve phytoplankton composition and abundance measurements that are as accurate as traditional microscopic analysis [3, 15]. However, FlowCAM systems require careful instrumental and experimental setups to yield reliable data [1]. Determining optimal settings often depends on *a priori* knowledge of particle density and sample composition [9, 1]. As a result, FlowCAM is more frequently used in controlled experiments and routine applications [8, 19, 21] than in exploratory surveys of less-studied systems, where its performance may be suboptimal [4, 9]. An additional and major challenge when using FlowCAM to analyse natural phytoplankton communities is the abundance of detritus particles, which can match or exceed phytoplankton particle counts [22, 2, 6]. While high abundance of detritus can hinder phytoplankton analysis in some applications, detritus itself can be the primary research focus in others. For example, detritus is an important factor in coastal seas that can be studied with FlowCAM systems [13]. Here, I present that a simple classifier based on logistic regression can be easily derived and applied with acceptable accuracy to a quick sorting of phytoplankton/detritus particles.

Logistic regression also identifies the most important features for distinguishing detritus particles (Table 2). In general, detritus particles present a more variable texture (higher SIGMAI), less compact and more elongated shape (smaller COMP and higher ELON). Surprisingly, optical features such as peak fluorescence in the red and yellow wavebands (CH1 and CH2, respectively) do not rank among the most important features. This is due to the experimental settings, specifically the fluorescence-triggered mode, which does not capture particles with low chlorophyll content [9, 5].

The logistic-regression-based classifier can reconstruct the size spectra of phytoplankton and detritus particles of natural samples. The quality of these reconstructions is conditioned to the classifier’s precision. While the aggregated spectra of all samples are well represented (Fig. 1), the analyses for the date-specific spectra are less accurate (Table 3 and Fig. SM1). Although this can be due to low particle counts in the smaller and larger size-classes (Table SM1), systematic biases in the particle properties may be responsible for the frequent misclassification of the more abundant middle-sized particles. For example, detritus particles can have different features forming subclasses, e.g., needlevs. rounded-shaped detritus [13], which could not be perfectly captured by a single classifier. This unaccounted diversity complicates using simple binary classifiers, and more comprehensive approaches could be necessary.

## 5 Conclusion

The use of simple binary classifiers applied to FlowCAM data offers significant potential for analysing particle size spectra of natural samples. While logistic regression can achieve reliable particle classification, further refinement is needed to improve accuracy, particularly in diverse and dynamic environments.

## Data accessibility

New data was not generated in this study. The code for the calculations is available in https://github.com/ovgarol/detritus-logistic-classification.

## Authors’ contributions

O.G.-O.: conceptualization, data curation, formal and statistical analysis, writing-original draft, writing-review and editing.

## A Performance metrics

Performance metrics are based on the amount of true positives (TP, particles correctly classified as detritus), true negatives (TN, particles correctly classified as phytoplankton), false positives (FP, particles incorrectly classified as detritus), and false negatives (FN, particles incorrectly classified as phytoplankton). These four values are summarized in the confusion matrix. The four performance metric herein reported are:

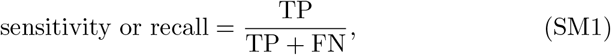

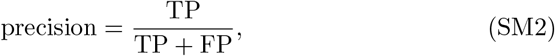

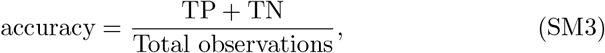

and

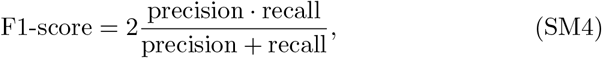

## B Additional figures

**Figure SM1:**
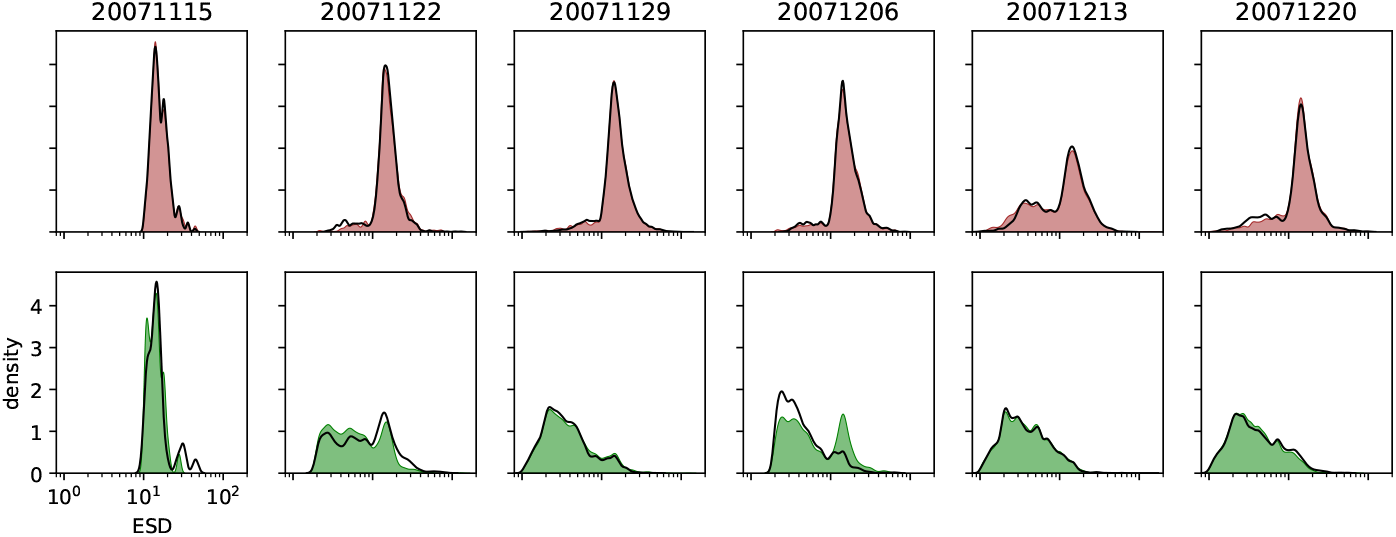
Observed (black line) and predicted size spectra (ESD, Equivalent Spherical diameter in μm) for detritus (brown area) and phytoplankton (green area) per sampling date.

**Table SM1:** Particle abundance per size-class and date.

| size-class ($\mu\text{m}$) | <2 | 2-5 | 5-10 | 10-20 | 20-50 | 50-100 | 100-200 |
| --- | --- | --- | --- | --- | --- | --- | --- |
| 20071115 | 0 | 0 | 0 | 285 | 54 | 0 | 0 |
| 20071122 | 10 | 512 | 421 | 1731 | 380 | 20 | 2 |
| 20071129 | 542 | 1699 | 778 | 3118 | 760 | 30 | 1 |
| 20071206 | 8 | 658 | 310 | 1921 | 551 | 22 | 0 |
| 20071213 | 806 | 2562 | 1414 | 1630 | 449 | 7 | 1 |
| 20071220 | 437 | 1225 | 628 | 1320 | 287 | 19 | 0 |

**Figure SM2:**
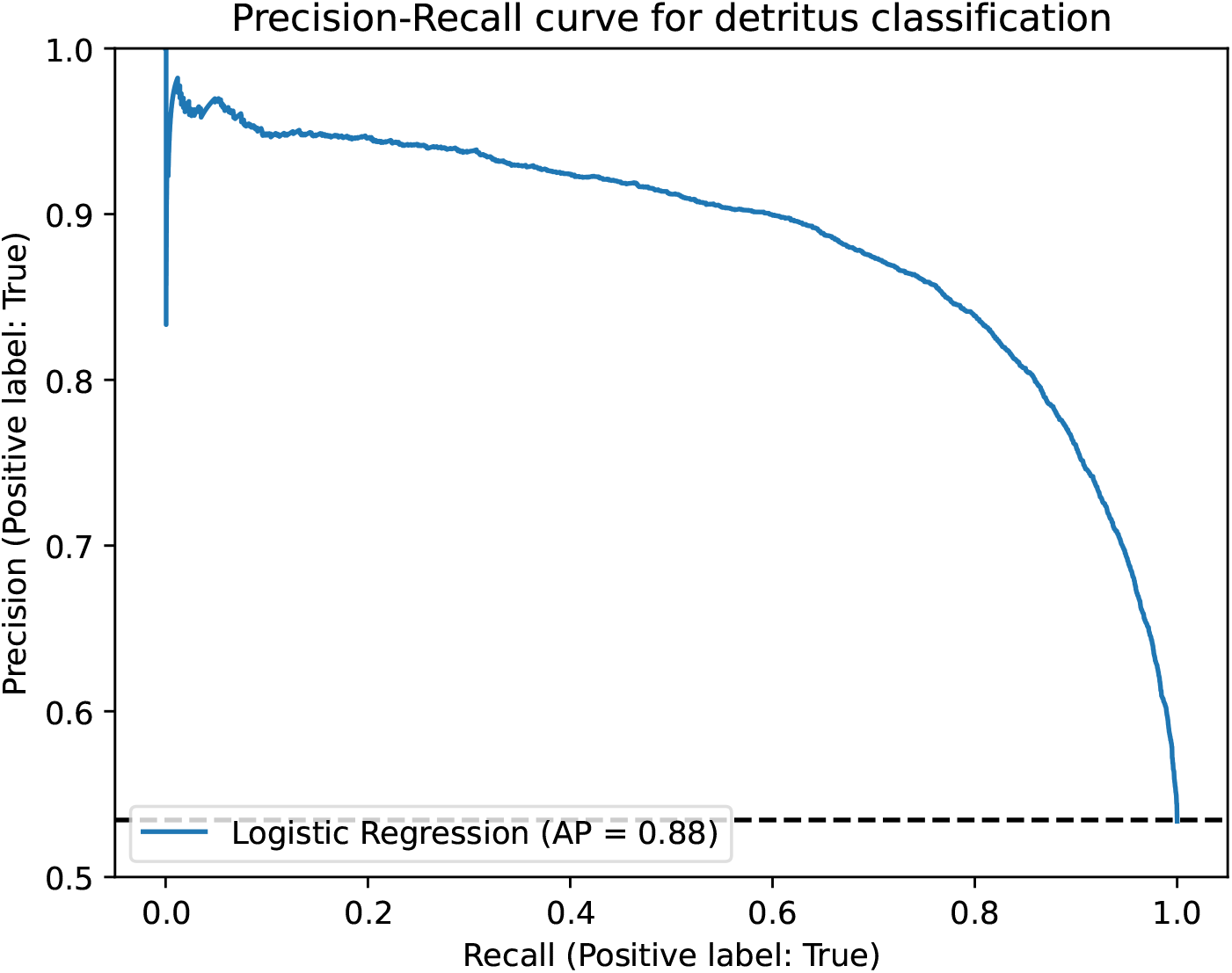
Precision-Recall curve for the model fitted with 30% of the dataset.

